# Restricted edition of ALS is required during co-edition in order to maintain normal tomato development

**DOI:** 10.1101/2025.08.27.672575

**Authors:** Kyoka Kuroiwa, Anaë Laurens, Marie-Noëlle Corre, Raphaël Lugan, Marianne Mazier, Jean-Luc Gallois

## Abstract

Co-editing strategies have emerged as an approach to facilitate the selection of CRISPR/Cas-mediated mutants in a transgene-free manner: the gene of interest is edited together with a reporter gene, whose mutation can be selected visually or pharmacologically. In this work, we asses the impact of editing the well-used reporter Acetolactate synthase (ALS) on plant development and metabolome. We show that the desired mutation in ALS1 (P186S) conferring selectable herbicide resistance trait does not show significant impact on the plant morphology and physiology but that the additional mutations resulting from the same sgRNA can result in reduced vegetative vigor and altered metabolomic profiles in tomato.

## Main text

Co-editing strategies have emerged as an approach to facilitate the selection of CRISPR/Cas-mediated mutants in a transgene-free manner: the gene of interest is edited together with a reporter gene, whose mutation can be selected visually or pharmacologically (Gallois and Nogué, 2023). *Acetolactate synthase* (*ALS*) is one of such reporter genes. *ALS* genes code for a key enzyme which is involved in the initial steps of branched-chain amino acid (BCAA) biosynthesis in chloroplast and targeted by major herbicide groups for controlling weeds. Eight nonsynonymous mutation sites have been found among the natural *ALS* allelic diversity in weed species and associated with dominant herbicide resistance (Yu and Powles, 2014). Recently, a genome-editing reporter system targeting one of those sites, the Proline-197 (in Arabidopsis thaliana *ALS*) equivalent codon, by a cytosine base editor (CBE) has been successfully implemented in tomato, tobacco, potato, and citrus *ALS1* genes, implying the wide applicability of this strategy in different crop species (Huang *et al*., 2023; Kuroiwa *et al*., 2023). However, Veillet *et al*. (2019) reported that the single guide RNA (sgRNA) commonly used to modify the targeted proline could trigger an additional nonsynonymous mutation in tomato and potato *ALS1* and in the highly homologous *ALS2* gene. Given *ALS*’s significant role in BCAA biosynthesis, we sought to ensure that extensive *ALS* editing would not impact the plant development and vigor.

For this objective, we assessed a panel of seven cherry tomato genotypes derived from previously obtained *SlALS*-edited population for their development and metabolomic profiles. The panel consisted of progenies of the *SlALS*-edited (Veillet *et al*., 2019) and the *SlALS*-*SleIF4E1*-co-edited (Kuroiwa *et al*., 2023) cherry tomato populations that were generated by CBE, both with the above-mentioned sgRNA targeting the *Sl*ALS1-P186 codon (equivalent of *At*ALS P197) (Figure 1a). The seven genotypes ranged from desired mutants harboring a single amino acid substitution at P186 in *Sl*ALS1 to those with two substitutions within the editing window of target sequence in *Sl*ALS1—at Q184 and P186—plus one substitution at Q182 in *Sl*ALS2 (*Sl*ALS1-Q184 equivalent) (Figure 1b, Table S1). We then subjected these plants to a whole-plant phenotyping and metabolomics study.

**Figure 1.**
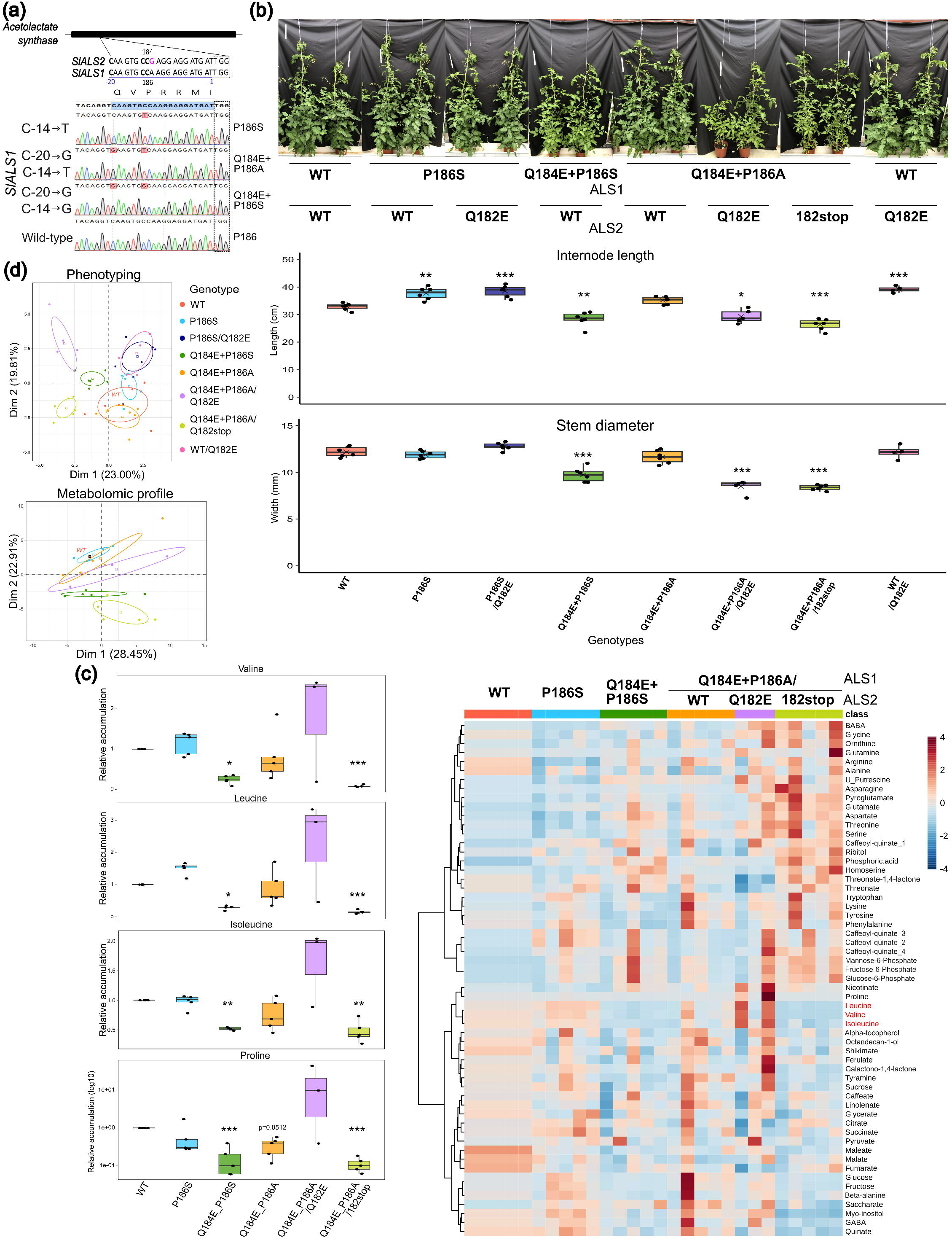
Cherry tomatoes with extensive base edition in one or more *ALS* genes show differential development and metabolomic profiles compared to the wild type. (a) Genomic region in *SlALS1* targeted for co-editing and the highly homologous region in *SlALS2* and the resulting editions. The sgRNA nucleotide sequence is underlined in blue and numbered according to the positions from the PAM site (dotted rectangle). The target codon *SlALS1*-P186 is indicated by “186” below and its equivalent *SlALS2*-P184 by “184”. Cytosines in the edition window are bolded. The expected amino acid substitutions due to the editing are on the right. (b) Morphology (top) and phenotypic measurements (bottom) at 45 days post transplanting of *ALS*-edited tomato plants. Genotypes are designated by the respective amino acid substitutions in *ALS1* and *ALS2*. Six plants per mutant genotype were measured except for *ALS1*^*Q184E+P186A*^*/ALS2*^*Q182E*^ (n=5) and *ALS2*^*Q182E*^ (n=4). (c) Right: Heatmap of primary metabolite accumulation in five *ALS*-edited mutants with normalization by wild type in each injection batch. Scale: from blue (minimum) to red (maximum peak area) within each metabolite class. Left: relative accumulation levels (peak areas) of BCAAs and proline, a stress-associated compound, in each genotype. Five plants per genotype were analyzed except for *ALS1*^*Q184E+P186A*^*/ALS2*^*Q182E*^ (n=3). (d) Principal component analysis individual plots based on the phenotypic measurements and metabolomic profiles. Ellipses represent 95% confidence interval for each genotype. Statistical significance of difference between each genotype and the WT was evaluated for (b) and (c) based on ANOVA followed by Dunnett’s test or Kruskal-Wallis test followed by Conover’s many-to-one comparison test and shown as asterisks p <0.05 *, <0.01 **, and <0.001 ***.

The *ALS*-edited and wild-type (WT) cherry tomato plants were grown for phenotypic assessment under greenhouse conditions in two independent sets, each containing six of the edited genotypes and the WT. In addition, the *ALS1*^*Q184E+P186A*^*/ALS2*^*Q182stop*^ genotype was included in one set due to its availability (Table S1, Figure S1). In the vegetative growth aspects, the plants harboring only one substitution at *Sl*ALS1-P186 showed similar or enhanced growth compared to the WT, regardless of the *ALS2* allele (*ALS1*^*P186S*^ and *ALS1*^*P186S*^*/ALS2*^*Q182E*^) as well as the plants with Q182E substitution in *SlALS2* alone (Figure 1b, S2, S3). However, the plants with additional Q184E substitution in *SlALS1* exhibited differential growth profiles depending on the P186 substitution: *ALS1*^*Q184E+P186S*^ showed reduced primary and secondary growth, while *ALS1*^*Q184E+P186A*^ showed normal to reduced growth with a large intra-genotypic variability (Figure 1b, S3). Meanwhile, the plants with three mutations in *ALS1* and *ALS2* combined, namely *ALS1*^*Q184E+P186A*^*/ALS2*^*Q182E*^ and *ALS1*^*Q184E+P186A*^*/ALS2*^*Q182stop*^, consistently showed reduced vegetative growth and lower leaf chlorophyll and nitrogen contents (Figure 1b, S2, S3). Nevertheless, edited genotypes did not show consistent differences from the WT in their reproductive growth across independent sets (Figure S4). Altogether, while the desired mutation by co-editing at *Sl*ALS1-P186 has no obvious impact on tomato growth, additional mutations in *Sl*ALS1 and/or *Sl*ALS2 inflict negative impact on the vegetative vigor.

Although the plants with *ALS1*^*P186S*^ allele did not show detectable difference from the WT in greenhouse conditions, their metabolic pathways involving BCAA could be affected, possibly leading to altered stress response. Therefore, we performed GC-EI-TOFMS profiling on the primary metabolites in leaves from the WT and five mutant genotypes grown in a climatic chamber (Table S1). The heatmap of 57 identified compounds revealed that the WT and the *ALS1*^*P186S*^ plants are largely similar in their primary metabolite accumulation patterns, including of BCAA and stress response-related compounds such as proline, GABA, and glutamate (Figure 1c, S7). In contrast, genotypes with reduced vegetative growth: *ALS1*^*Q184E+P186S*^ and *ALS1*^*Q184E+P186A*^*/ALS2*^*Q182stop*^ exhibited an opposite trend of accumulations across identified compounds such as phosphoric acid and homoserine and significantly lower BCAA and proline accumulation levels (Figure 1c). Notably, the metabolomic profiles of the genotypes *ALS1*^*Q184E+P186A*^ and *ALS1*^*Q184E+P186A*^*/ALS2*^*Q182E*^ were highly variable between individuals (Figure 1c). Finally, the principal component analysis indicates that the growth and metabolomic profiles of the ALS mutants are clustered in a similar manner, supporting that the *SlALS1*-P186S mutation maintains the WT-like phenotypes, while the excessive *ALS1* and *ALS2* mutations induce changes in the metabolomic profiles, leading to alteration of plant development (Figure 1d).

In summary, we analyzed the developmental and metabolomic outcomes of editing *ALS* in the cherry tomato using various mutant lines obtained with a commonly used *ALS1*-targeting sgRNA. We show that the desired mutation in *ALS1* (P186S) conferring selectable herbicide resistance trait does not show significant impact on the plant morphology and physiology but that the additional mutations resulting from the same sgRNA can result in reduced vegetative vigor and altered metabolomic profiles in tomato. Present work validates the advantage of the co-editing strategy targeting the *ALS*, while also warning that the strategy must be used with careful choice of the plant transformation/selection protocol such as transient expression of the base editor cassette (Huang *et al*., 2023; Veillet *et al*., 2019) and with alternative tools like the high precision CBE (Tan *et al*., 2019) or the prime editors (Vu *et al*., 2023) in order to avoid the risks of generating complex *ALS* mutants. An eventual reversion of the edited *ALS* gene can be also considered if the backcrossing of the edited plants are not suitable due to their mode of propagation or the genetic linkage between the target gene and the *ALS* loci (Gallois and Nogué, 2023; Guyon-Debast *et al*., 2021). With this work, we hope to provide the guidelines and precautions for more efficient use of ALS co-editing strategy in future experiments.

## Acknowledgement

The authors thank Emmanuel Botton (INRAE GAFL) for taking care of tomato plants throughout the experiments conducted for this work, Mathilde Causse (INRAE GAFL) for her advice regarding the generation of the panel of genotypes in this study, and Robin Thibert (INRAE GAFL) for acquisition of the preliminary data. The authors also thank Fabien Nogué (INRAE Versailles IJPB) and Florian Veillet (CEA Cadarache) for their valuable insights during the article writing.

## Supporting Information

**Table S1** Panel of ALS mutants used in this work;

**Figure S1** Experimental setup of the whole-plant phenotyping in the greenhouse;

**Figure S2** Mutations outside of ALS1 P186 site affect vegetative growth of cherry tomato in set B;

**Figure S3** ALS mutations show differential effects on additional vegetative parameters.

**Figure S4** ALS mutations do not show genotype-dependent effects on phenological trait.

**Figure S5** *ALS* mutations do not show genotype-dependent effects on reproductive traits.

**Figure S6** ALS mutations do not show genotype-dependent effects on tomato yield.

**Figure S7** Detailed metabolomic profiles of *ALS* mutant cherry tomatoes.

**Figure S8** Principal component analysis (PCA) variable plots on the developmental measurements and the metabolomic profiles.

**Figure S9** Principal component analysis (PCA) on the developmental measurements of set B and the sets A and B combined.

**Data S1** List of 57 metabolites identified in GC-EI-TOFMS

## Notes

### Competing Interest Statement

The authors have declared no competing interest.

