## Supplementary material for "Restricted edition of ALS is required during co-edition in order to maintain normal tomato development": sup data

### Slide 1
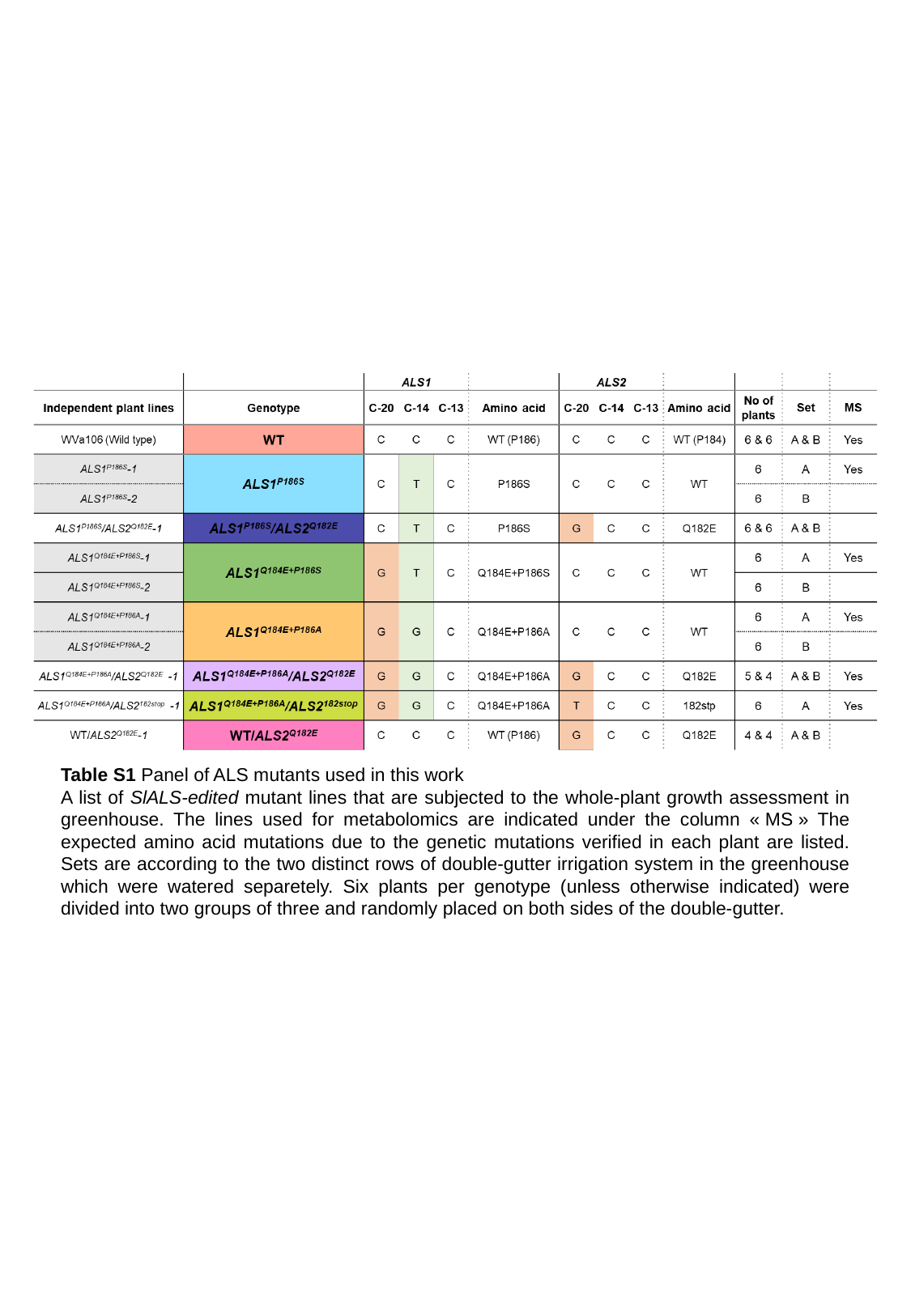

Table S1 Panel of ALS mutants used in this work
A list of SlALS-edited mutant lines that are subjected to the whole-plant growth assessment in greenhouse. The lines used for metabolomics are indicated under the column « MS » The expected amino acid mutations due to the genetic mutations verified in each plant are listed. Sets are according to the two distinct rows of double-gutter irrigation system in the greenhouse which were watered separetely. Six plants per genotype (unless otherwise indicated) were divided into two groups of three and randomly placed on both sides of the double-gutter.

### Slide 2
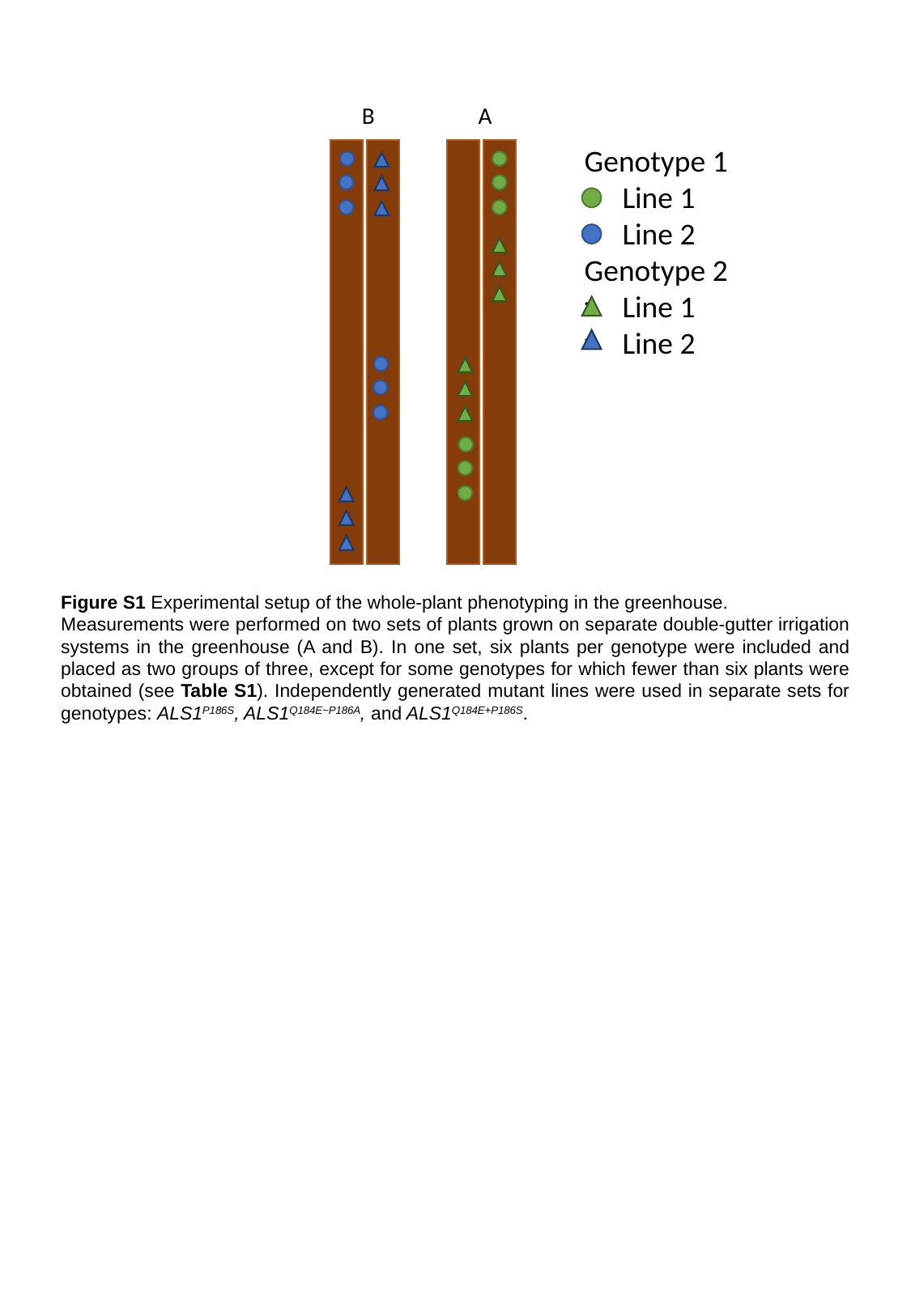

B
A
Genotype 1
Line 1
Line 2
Genotype 2
Line 1
Line 2
Figure S1 Experimental setup of the whole-plant phenotyping in the greenhouse.
Measurements were performed on two sets of plants grown on separate double-gutter irrigation systems in the greenhouse (A and B). In one set, six plants per genotype were included and placed as two groups of three, except for some genotypes for which fewer than six plants were obtained (see Table S1). Independently generated mutant lines were used in separate sets for genotypes: ALS1^P186S^, ALS1^Q184E~P186A^, and ALS1^Q184E+P186S^.

### Slide 3
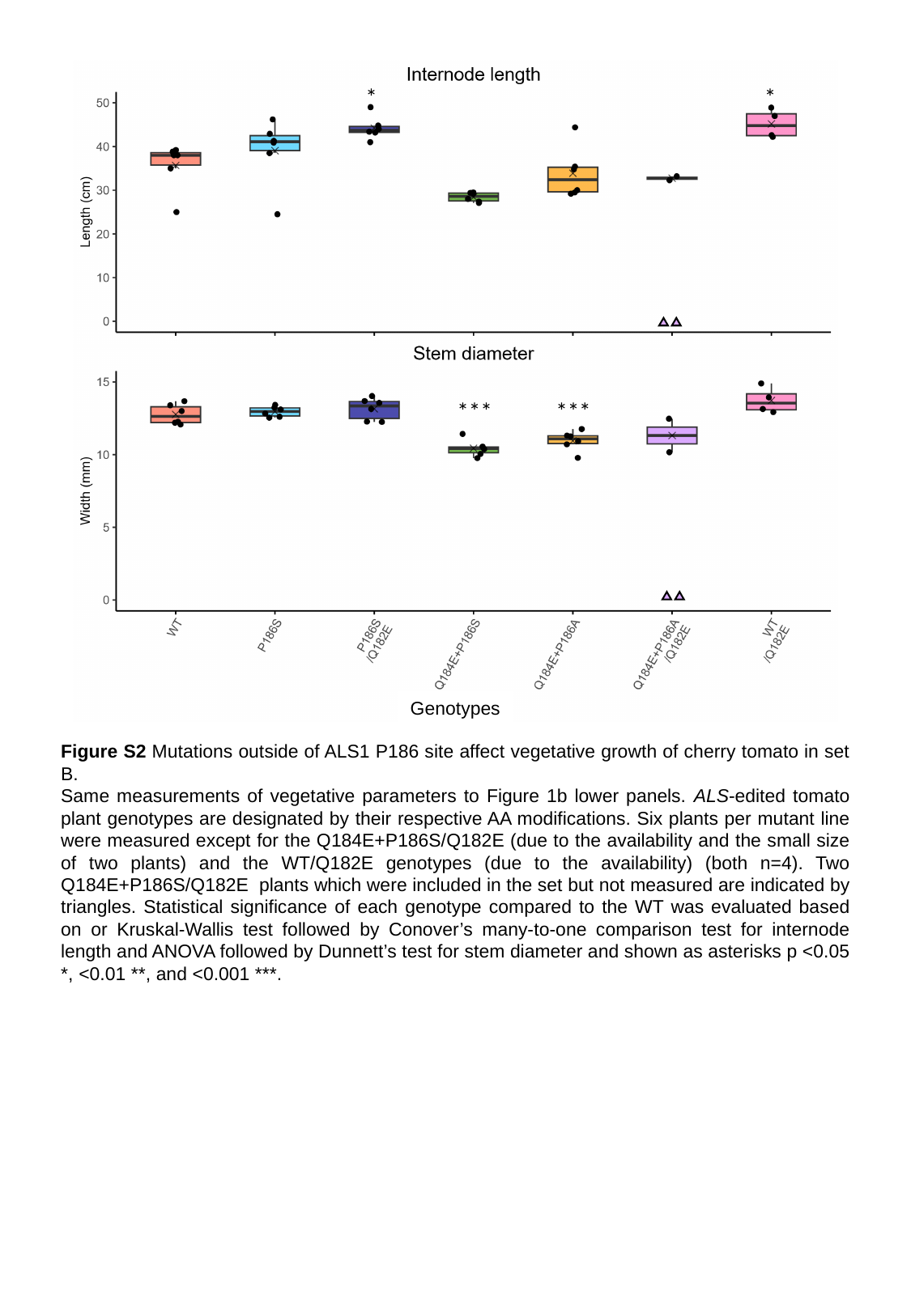

*
*
***
***
Genotypes
Figure S2 Mutations outside of ALS1 P186 site affect vegetative growth of cherry tomato in set B.
Same measurements of vegetative parameters to Figure 1b lower panels. ALS-edited tomato plant genotypes are designated by their respective AA modifications. Six plants per mutant line were measured except for the Q184E+P186S/Q182E (due to the availability and the small size of two plants) and the WT/Q182E genotypes (due to the availability) (both n=4). Two Q184E+P186S/Q182E plants which were included in the set but not measured are indicated by triangles. Statistical significance of each genotype compared to the WT was evaluated based on or Kruskal-Wallis test followed by Conover’s many-to-one comparison test for internode length and ANOVA followed by Dunnett’s test for stem diameter and shown as asterisks p <0.05 *, <0.01 **, and <0.001 ***.

### Slide 4
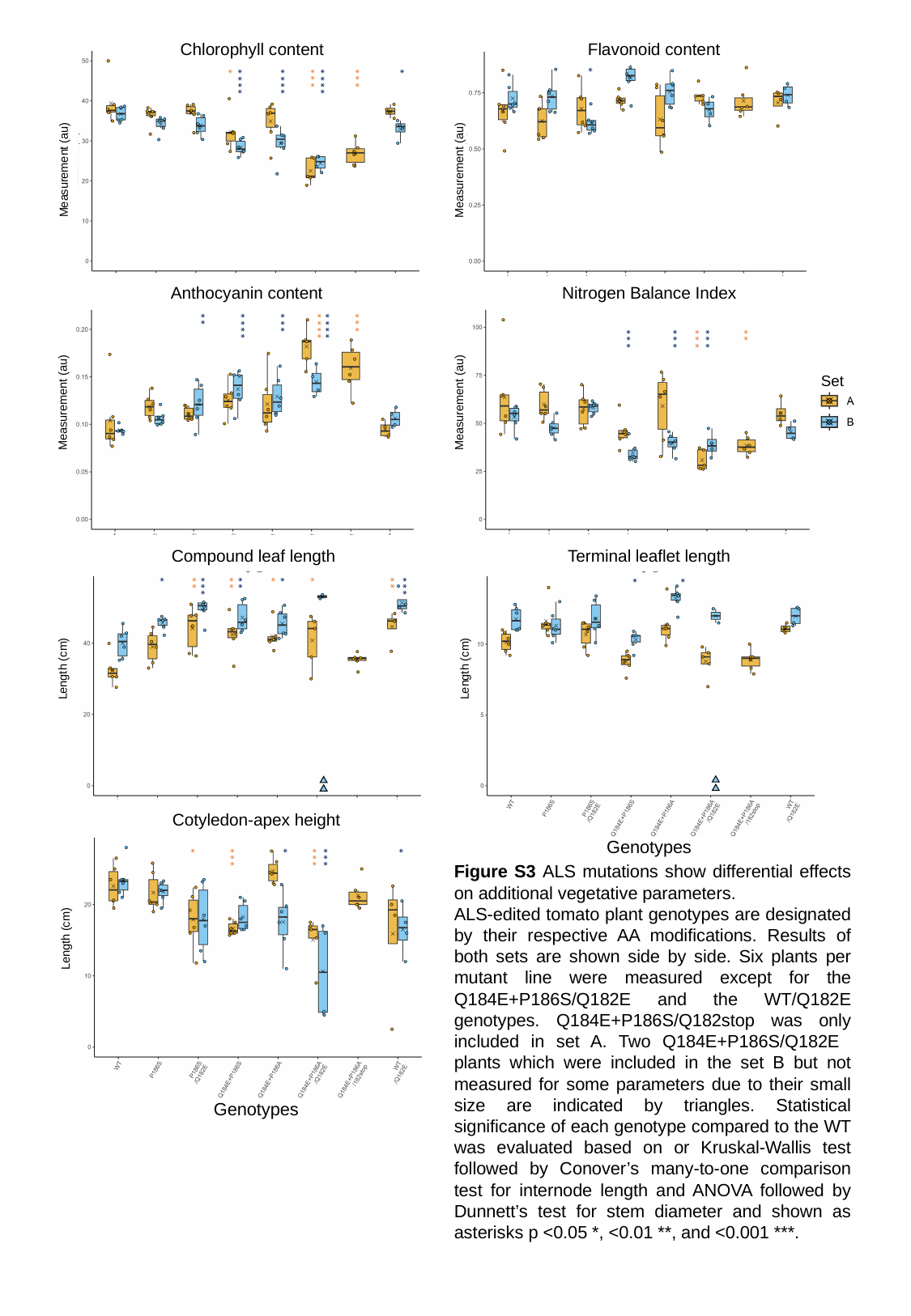

Chlorophyll content
*
*
***
***
****
****
****
Flavonoid content
*
Measurement (au)
Measurement (au)
Anthocyanin content
Nitrogen Balance Index
**
***
***
****
****
****
**
***
***
***
***
Set
Measurement (au)
Measurement (au)
Compound leaf length
Terminal leaflet length
*
*
*
*
**
**
**
**
***
***
*
*
Length (cm)
Length (cm)
Cotyledon-apex height
*
*
*
***
***
***
Genotypes
Figure S3 ALS mutations show differential effects on additional vegetative parameters.
ALS-edited tomato plant genotypes are designated by their respective AA modifications. Results of both sets are shown side by side. Six plants per mutant line were measured except for the Q184E+P186S/Q182E and the WT/Q182E genotypes. Q184E+P186S/Q182stop was only included in set A. Two Q184E+P186S/Q182E plants which were included in the set B but not measured for some parameters due to their small size are indicated by triangles. Statistical significance of each genotype compared to the WT was evaluated based on or Kruskal-Wallis test followed by Conover’s many-to-one comparison test for internode length and ANOVA followed by Dunnett’s test for stem diameter and shown as asterisks p <0.05 *, <0.01 **, and <0.001 ***.
Length (cm)
Genotypes

### Slide 5
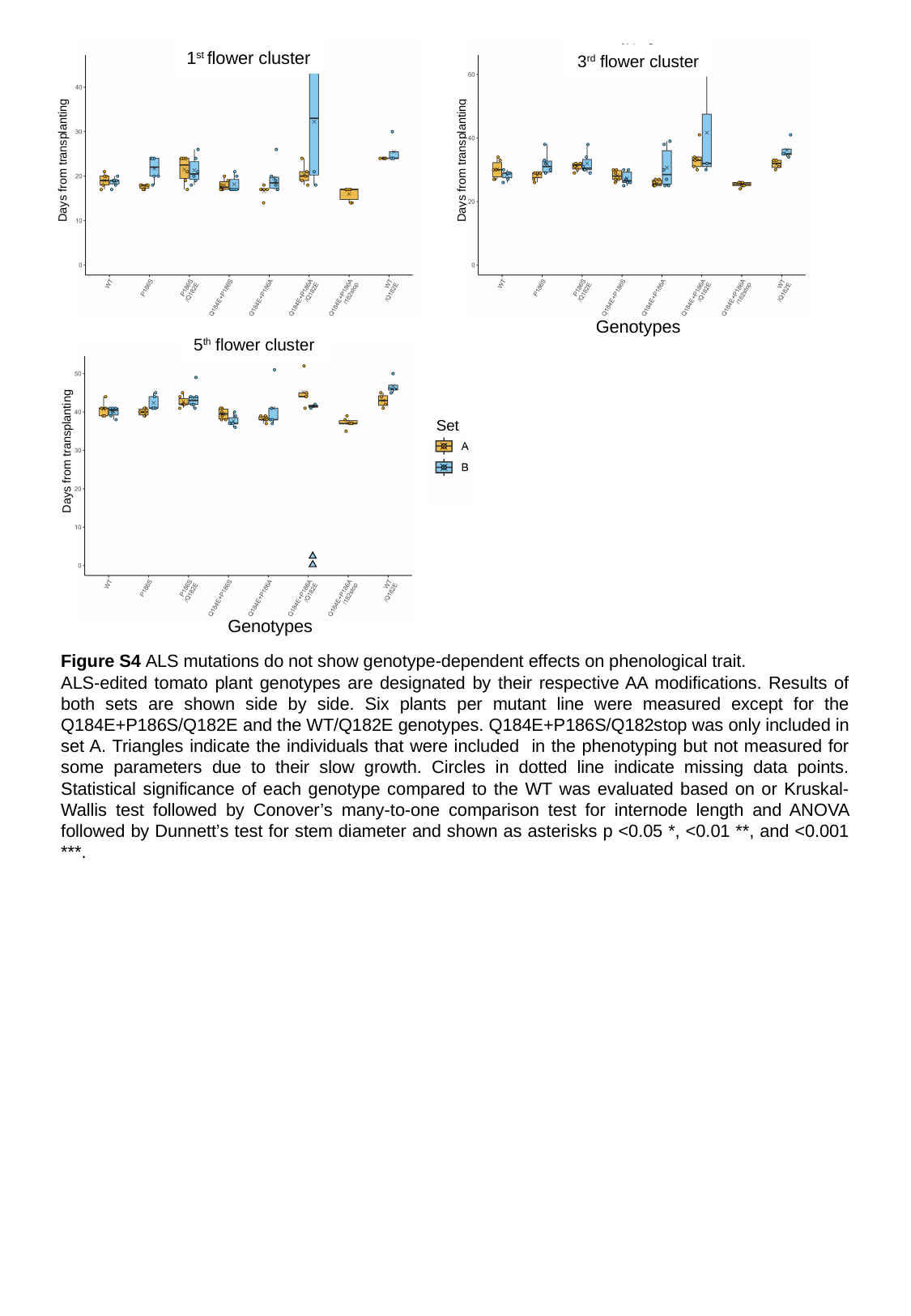

1^st^ flower cluster
3^rd^ flower cluster
Days from transplanting
Days from transplanting
Genotypes
5^th^ flower cluster
Days from transplanting
Set
Genotypes
Figure S4 ALS mutations do not show genotype-dependent effects on phenological trait.
ALS-edited tomato plant genotypes are designated by their respective AA modifications. Results of both sets are shown side by side. Six plants per mutant line were measured except for the Q184E+P186S/Q182E and the WT/Q182E genotypes. Q184E+P186S/Q182stop was only included in set A. Triangles indicate the individuals that were included in the phenotyping but not measured for some parameters due to their slow growth. Circles in dotted line indicate missing data points. Statistical significance of each genotype compared to the WT was evaluated based on or Kruskal-Wallis test followed by Conover’s many-to-one comparison test for internode length and ANOVA followed by Dunnett’s test for stem diameter and shown as asterisks p <0.05 *, <0.01 **, and <0.001 ***.

### Slide 6
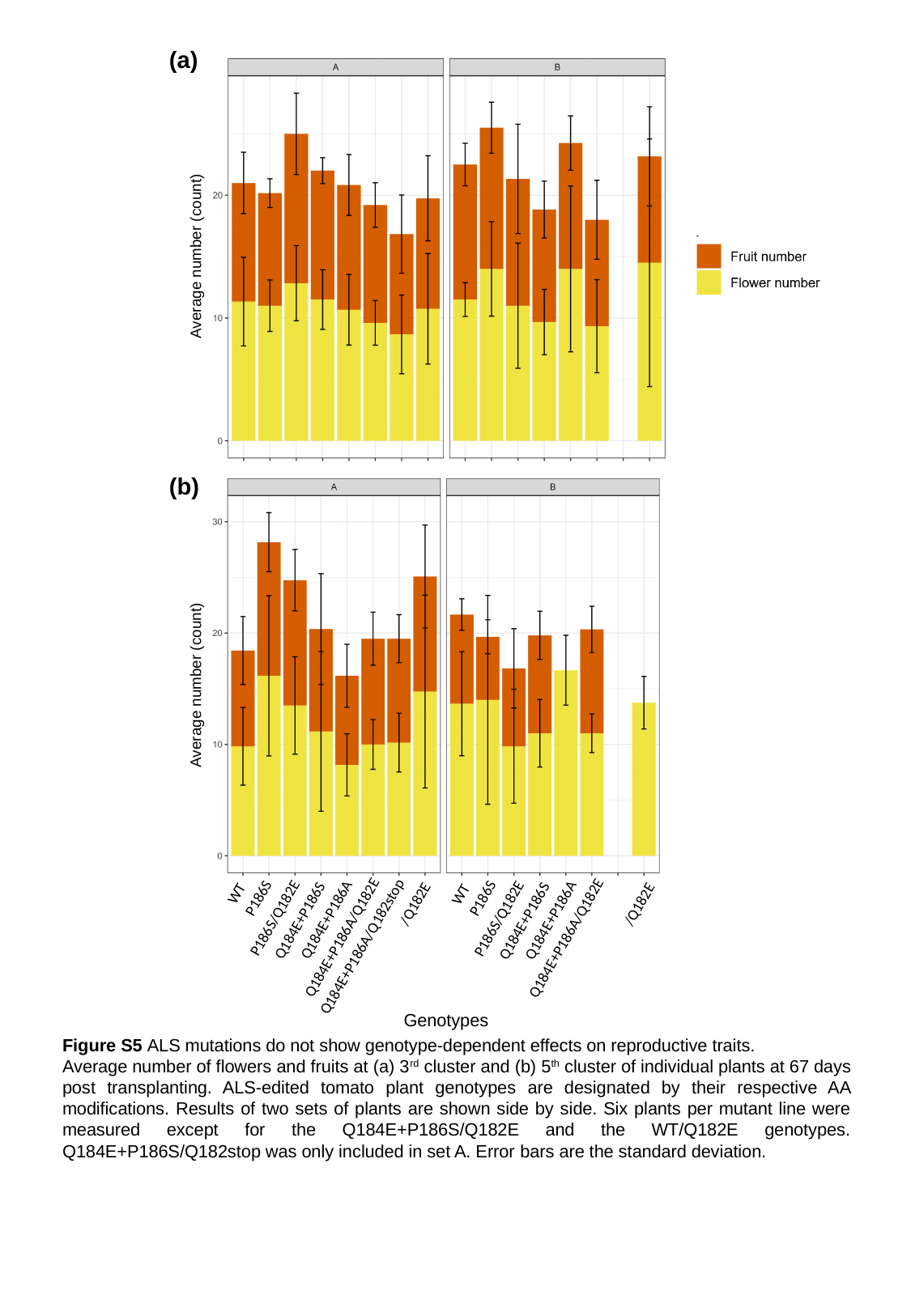

(a)
Average number (count)
(b)
Average number (count)
/Q182E
WT
/Q182E
WT
P186S
P186S
P186S/Q182E
P186S/Q182E
Q184E+P186A
Q184E+P186A
Q184E+P186S
Q184E+P186S
Q184E+P186A/Q182E
Q184E+P186A/Q182E
Q184E+P186A/Q182stop
Genotypes
Figure S5 ALS mutations do not show genotype-dependent effects on reproductive traits.
Average number of flowers and fruits at (a) 3^rd^ cluster and (b) 5^th^ cluster of individual plants at 67 days post transplanting. ALS-edited tomato plant genotypes are designated by their respective AA modifications. Results of two sets of plants are shown side by side. Six plants per mutant line were measured except for the Q184E+P186S/Q182E and the WT/Q182E genotypes. Q184E+P186S/Q182stop was only included in set A. Error bars are the standard deviation.

### Slide 7
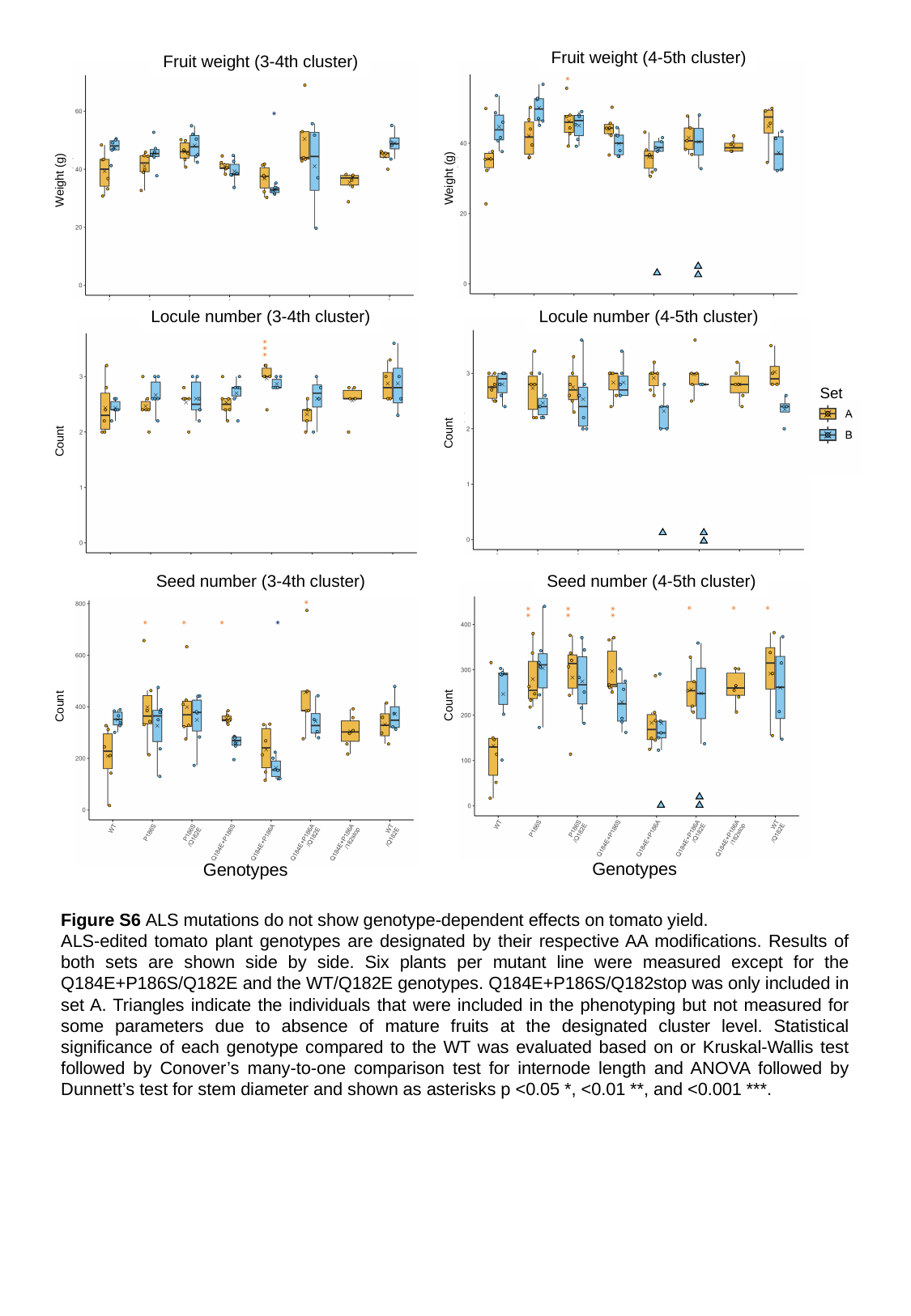

Fruit weight (4-5th cluster)
Fruit weight (3-4th cluster)
*
*
Weight (g)
Weight (g)
Locule number (3-4th cluster)
Locule number (4-5th cluster)
***
Set
Count
Count
Seed number (3-4th cluster)
Seed number (4-5th cluster)
*
*
*
**
**
**
*
*
*
*
*
Count
Count
Genotypes
Genotypes
Figure S6 ALS mutations do not show genotype-dependent effects on tomato yield.
ALS-edited tomato plant genotypes are designated by their respective AA modifications. Results of both sets are shown side by side. Six plants per mutant line were measured except for the Q184E+P186S/Q182E and the WT/Q182E genotypes. Q184E+P186S/Q182stop was only included in set A. Triangles indicate the individuals that were included in the phenotyping but not measured for some parameters due to absence of mature fruits at the designated cluster level. Statistical significance of each genotype compared to the WT was evaluated based on or Kruskal-Wallis test followed by Conover’s many-to-one comparison test for internode length and ANOVA followed by Dunnett’s test for stem diameter and shown as asterisks p <0.05 *, <0.01 **, and <0.001 ***.

### Slide 8
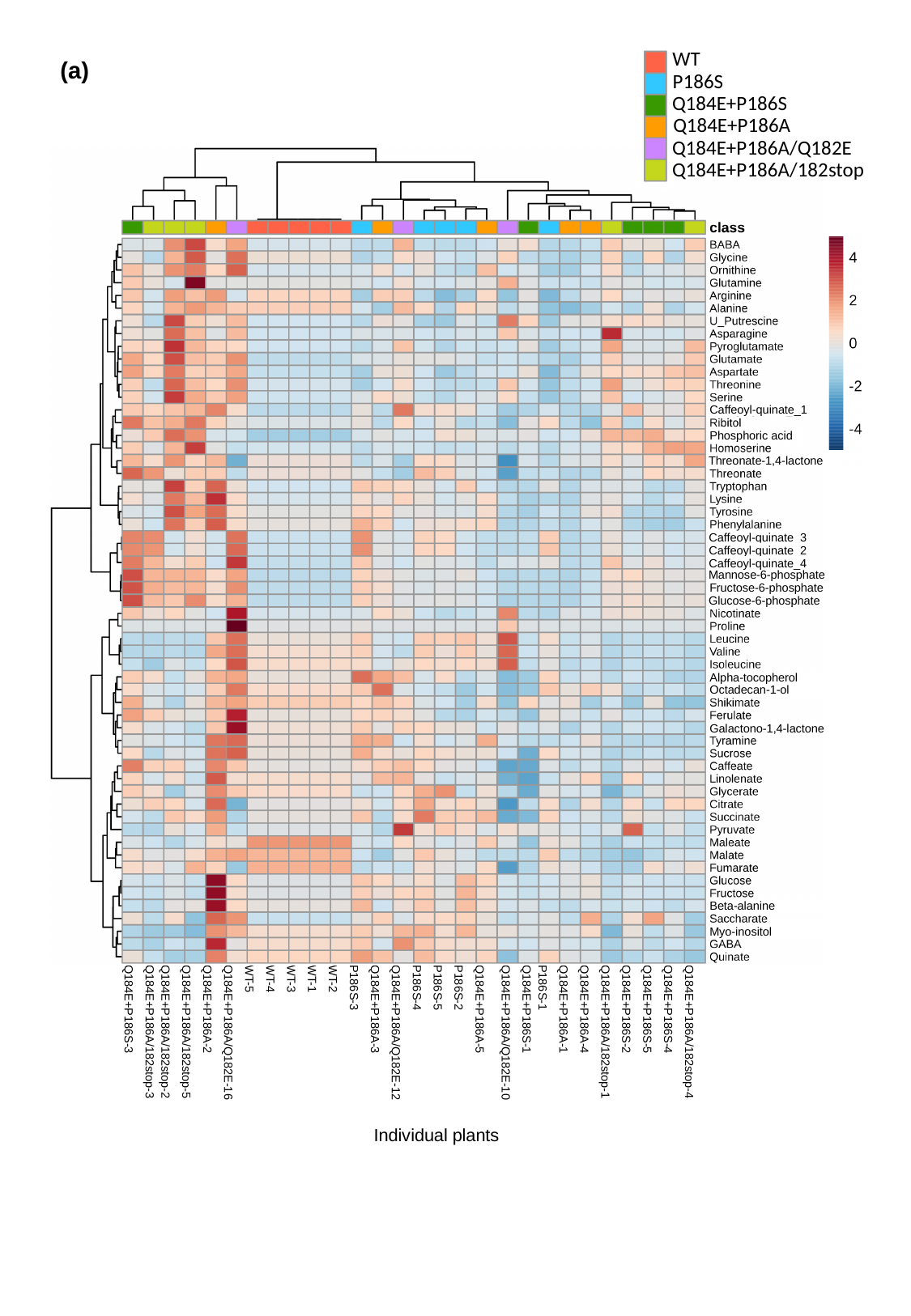

WT
P186S
Q184E+P186S
Q184E+P186A
Q184E+P186A/Q182E
Q184E+P186A/182stop
(a)
Caffeoyl-quinate_1
Phosphoric acid
Threonate-1,4-lactone
Caffeoyl-quinate_3
Caffeoyl-quinate_2
Caffeoyl-quinate_4
Mannose-6-phosphate
Fructose-6-phosphate
Glucose-6-phosphate
Alpha-tocopherol
Octadecan-1-ol
Galactono-1,4-lactone
Beta-alanine
Myo-inositol
| Q184E+P186S-3 | Q184E+P186A/182stop-3 | Q184E+P186A/182stop-2 | Q184E+P186A/182stop-5 | Q184E+P186A-2 | Q184E+P186A/Q182E-16 | WT-5 | WT-4 | WT-3 | WT-1 | WT-2 | P186S-3 | Q184E+P186A-3 | Q184E+P186A/Q182E-12 | P186S-4 | P186S-5 | P186S-2 | Q184E+P186A-5 | Q184E+P186A/Q182E-10 | Q184E+P186S-1 | P186S-1 | Q184E+P186A-1 | Q184E+P186A-4 | Q184E+P186A/182stop-1 | Q184E+P186S-2 | Q184E+P186S-5 | Q184E+P186S-4 | Q184E+P186A/182stop-4 |
| --- | --- | --- | --- | --- | --- | --- | --- | --- | --- | --- | --- | --- | --- | --- | --- | --- | --- | --- | --- | --- | --- | --- | --- | --- | --- | --- | --- |
Individual plants

### Slide 9
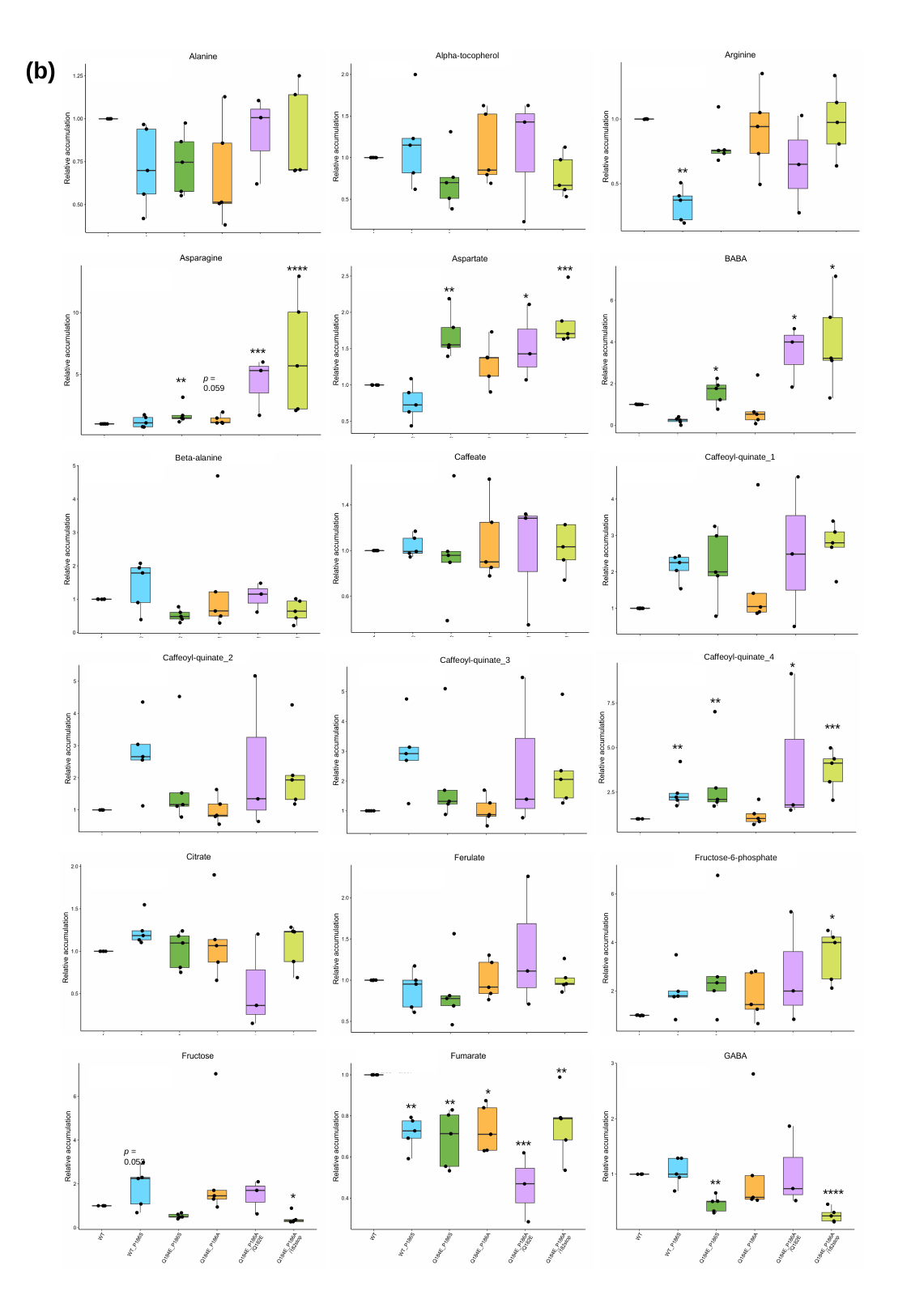

Alpha-tocopherol
(b)
p-value = 0.37
p-value = 0.133
p-value = 0.025
**
*
****
***
p-value = 0.001
p-value = 0.001
p-value = 0.001
**
*
*
***
*
p = 0.059
**
Caffeoyl-quinate_1
Beta-alanine
p-value = 0.151
p-value = 0.943
p-value = 0.11
Caffeoyl-quinate_4
Caffeoyl-quinate_2
Caffeoyl-quinate_3
*
p-value = 0.043
p-value = 0.005
p-value = 0.052
**
***
**
Fructose-6-phosphate
p-value = 0.527
p-value = 0.098
p-value = 0.277
*
p-value = 0.007
**
p-value = 0.002
p-value = 0.002
*
**
**
***
p = 0.052
**
****
*

### Slide 10
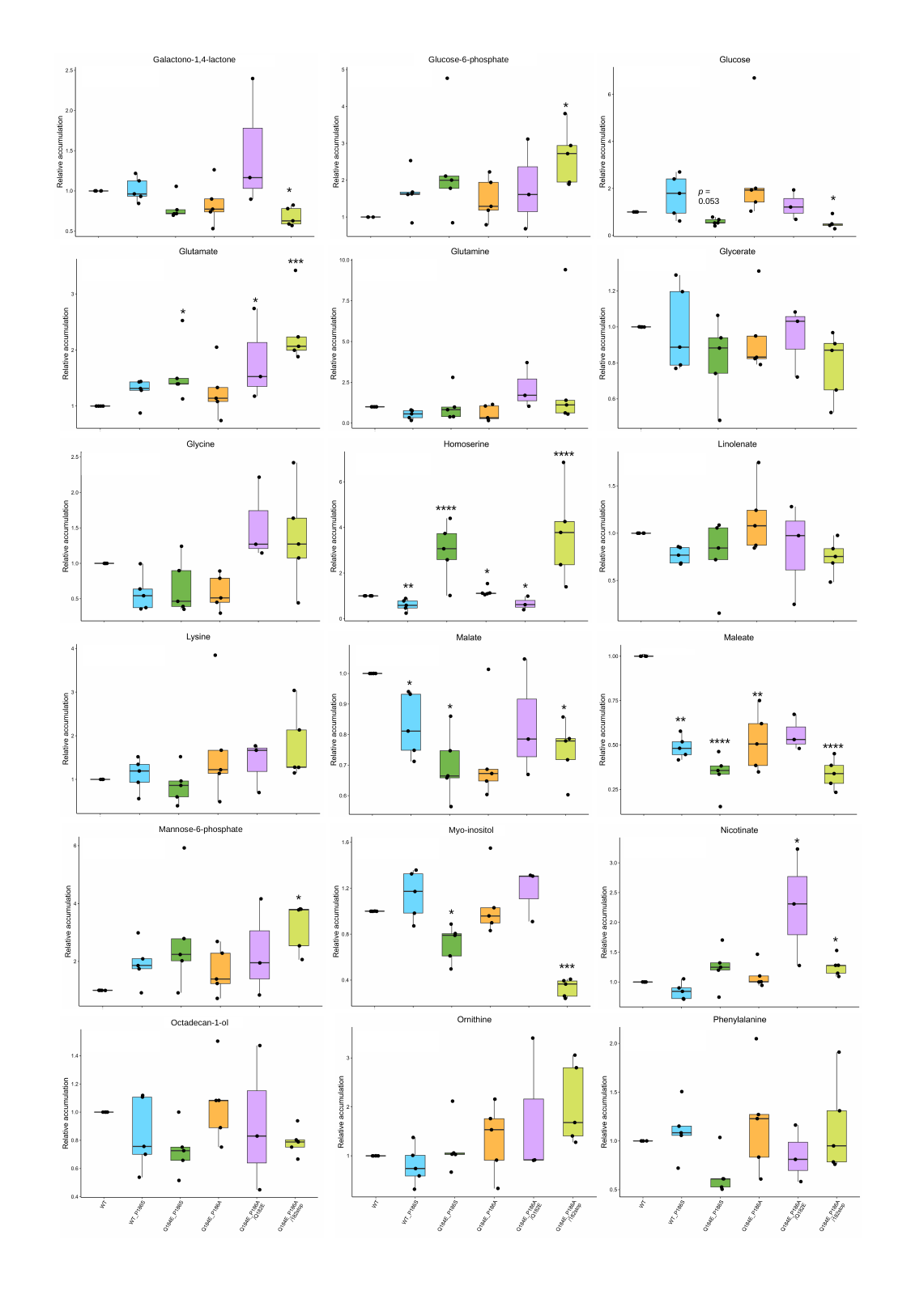

Galactono-1,4-lactone
Glucose-6-phosphate
p-value = 0.002
p-value = 0.02
p-value = 0.076
*
*
p = 0.053
*
***
p-value = 0.371
p-value = 0.007
p-value = 0.066
*
*
****
p-value = 0.01
p-value = 0.071
p-value < 10^-4^
****
*
**
*
p-value = 0.001
p-value = 0.152
p-value = 0.043
*
**
*
*
**
****
****
Mannose-6-phosphate
Myo-inositol
*
p-value = 0.006
p-value = 0.002
p-value = 0.089
*
*
*
***
Octadecan-1-ol
p-value = 0.178
p-value = 0.122
p-value = 0.15

### Slide 11
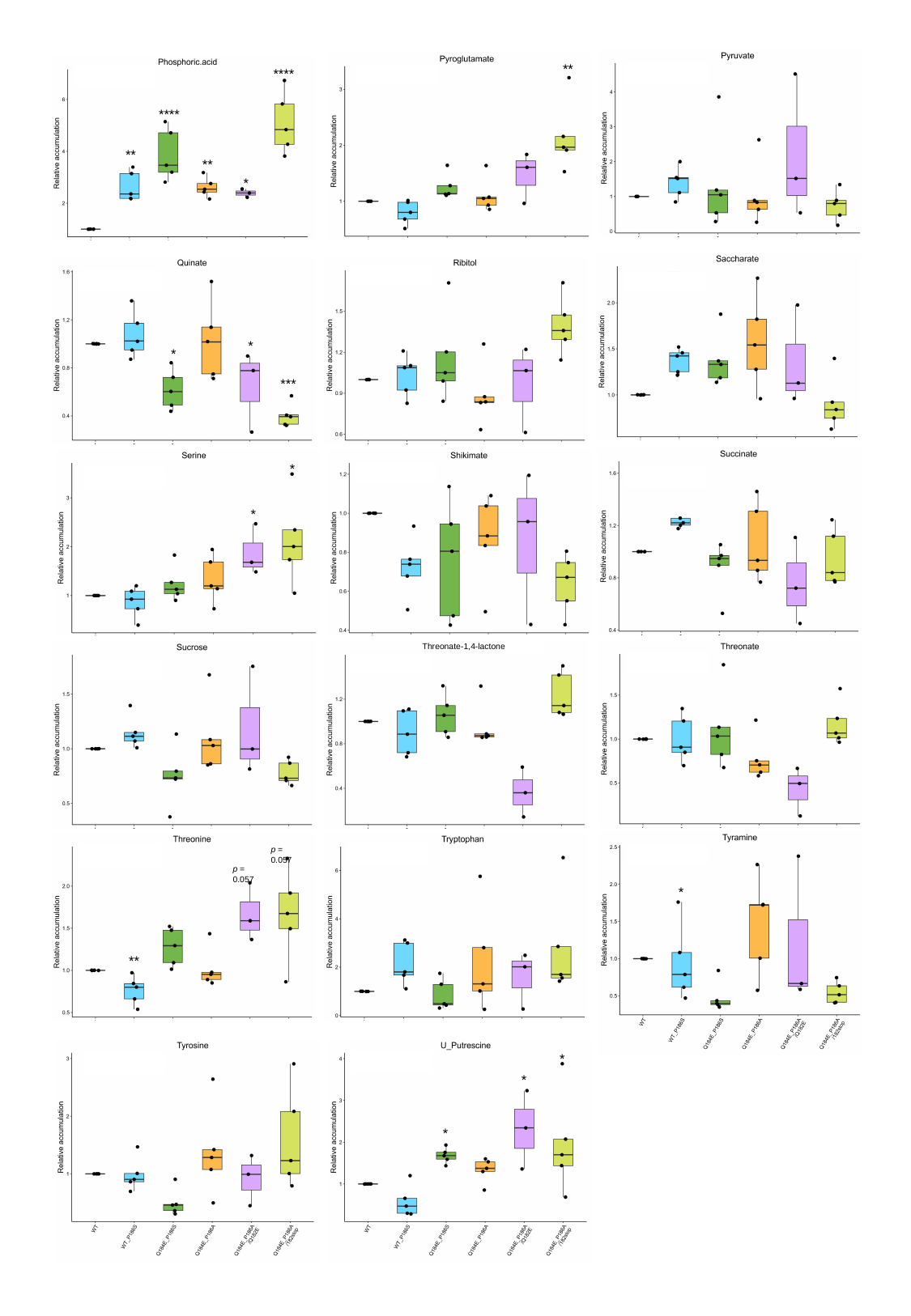

**
****
p-value = 0.015
p-value = 0.035
p-value = 0.015
****
**
**
*
p-value = 0.004
p-value = 0.035
p-value = 0.015
*
*
***
*
p-value = 0.004
p-value = 0.015
p-value = 0.035
*
Threonate-1,4-lactone
p-value = 0.015
p-value = 0.004
p-value = 0.035
p = 0.057
p-value = 0.004
p-value = 0.015
p-value = 0.035
p = 0.057
*
**
*
p-value = 0.004
p-value = 0.035
*
*

### Slide 12
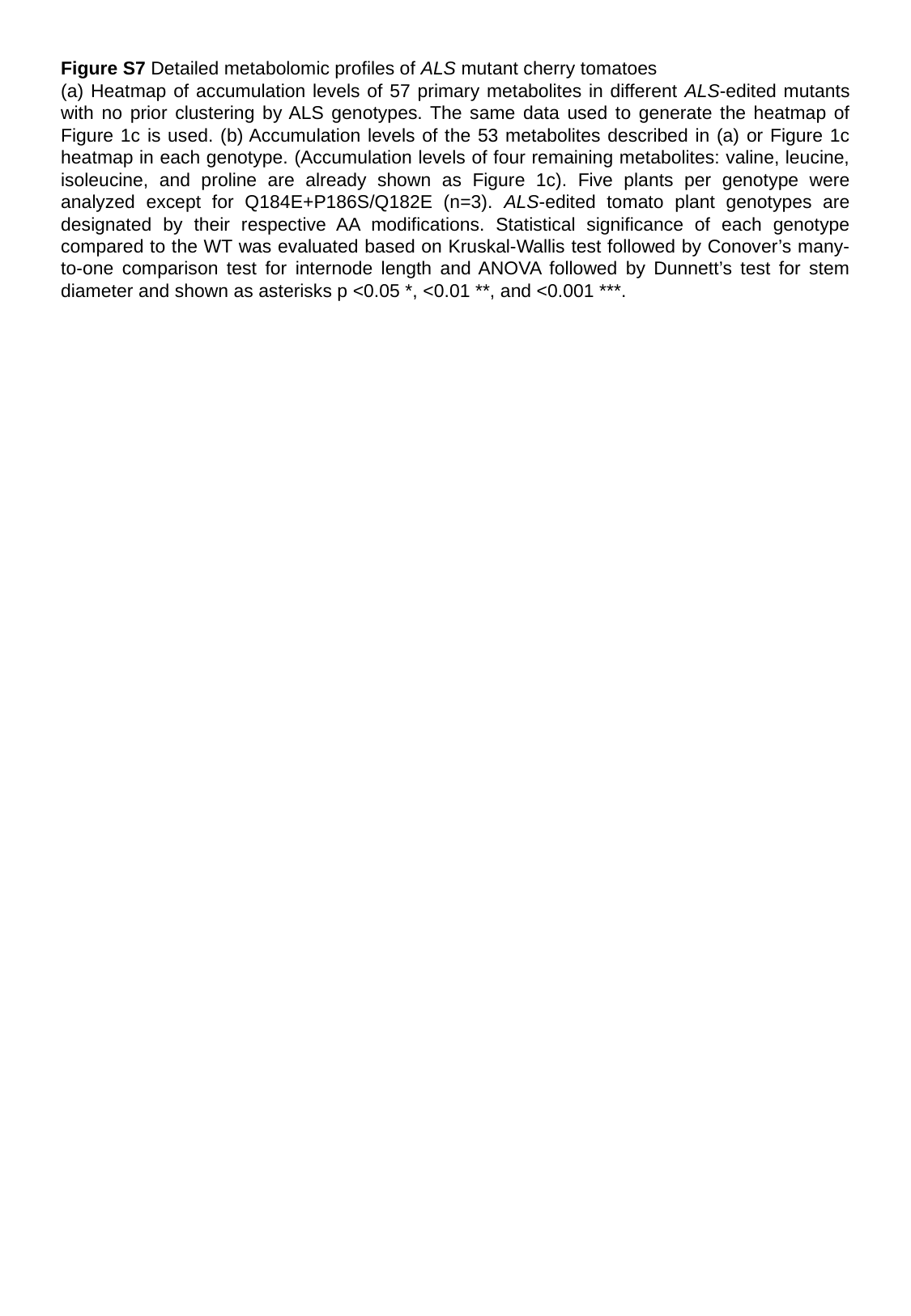

Figure S7 Detailed metabolomic profiles of ALS mutant cherry tomatoes
(a) Heatmap of accumulation levels of 57 primary metabolites in different ALS-edited mutants with no prior clustering by ALS genotypes. The same data used to generate the heatmap of Figure 1c is used. (b) Accumulation levels of the 53 metabolites described in (a) or Figure 1c heatmap in each genotype. (Accumulation levels of four remaining metabolites: valine, leucine, isoleucine, and proline are already shown as Figure 1c). Five plants per genotype were analyzed except for Q184E+P186S/Q182E (n=3). ALS-edited tomato plant genotypes are designated by their respective AA modifications. Statistical significance of each genotype compared to the WT was evaluated based on Kruskal-Wallis test followed by Conover’s many-to-one comparison test for internode length and ANOVA followed by Dunnett’s test for stem diameter and shown as asterisks p <0.05 *, <0.01 **, and <0.001 ***.

### Slide 13
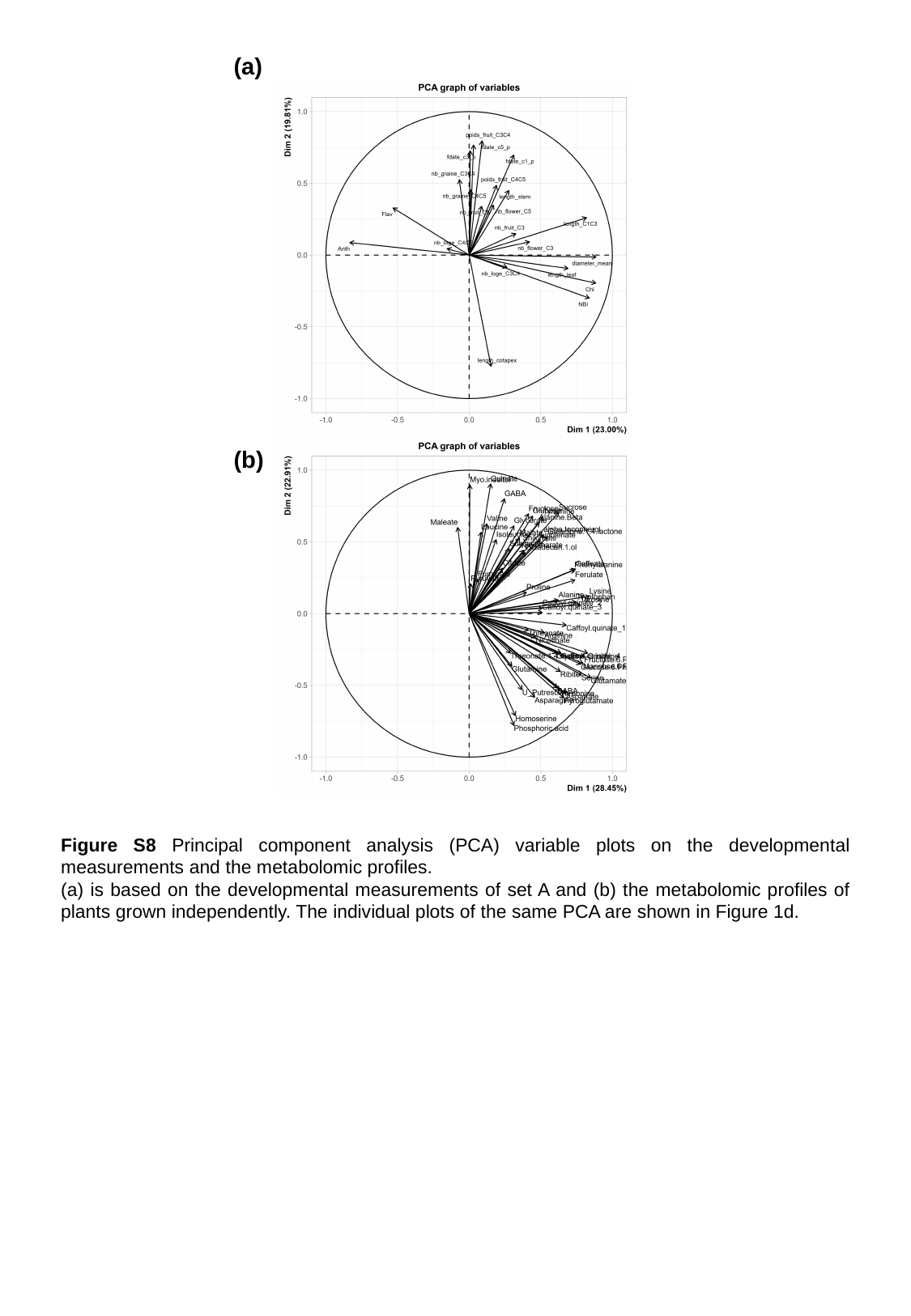

(a)
(b)
Figure S8 Principal component analysis (PCA) variable plots on the developmental measurements and the metabolomic profiles.
(a) is based on the developmental measurements of set A and (b) the metabolomic profiles of plants grown independently. The individual plots of the same PCA are shown in Figure 1d.

### Slide 14
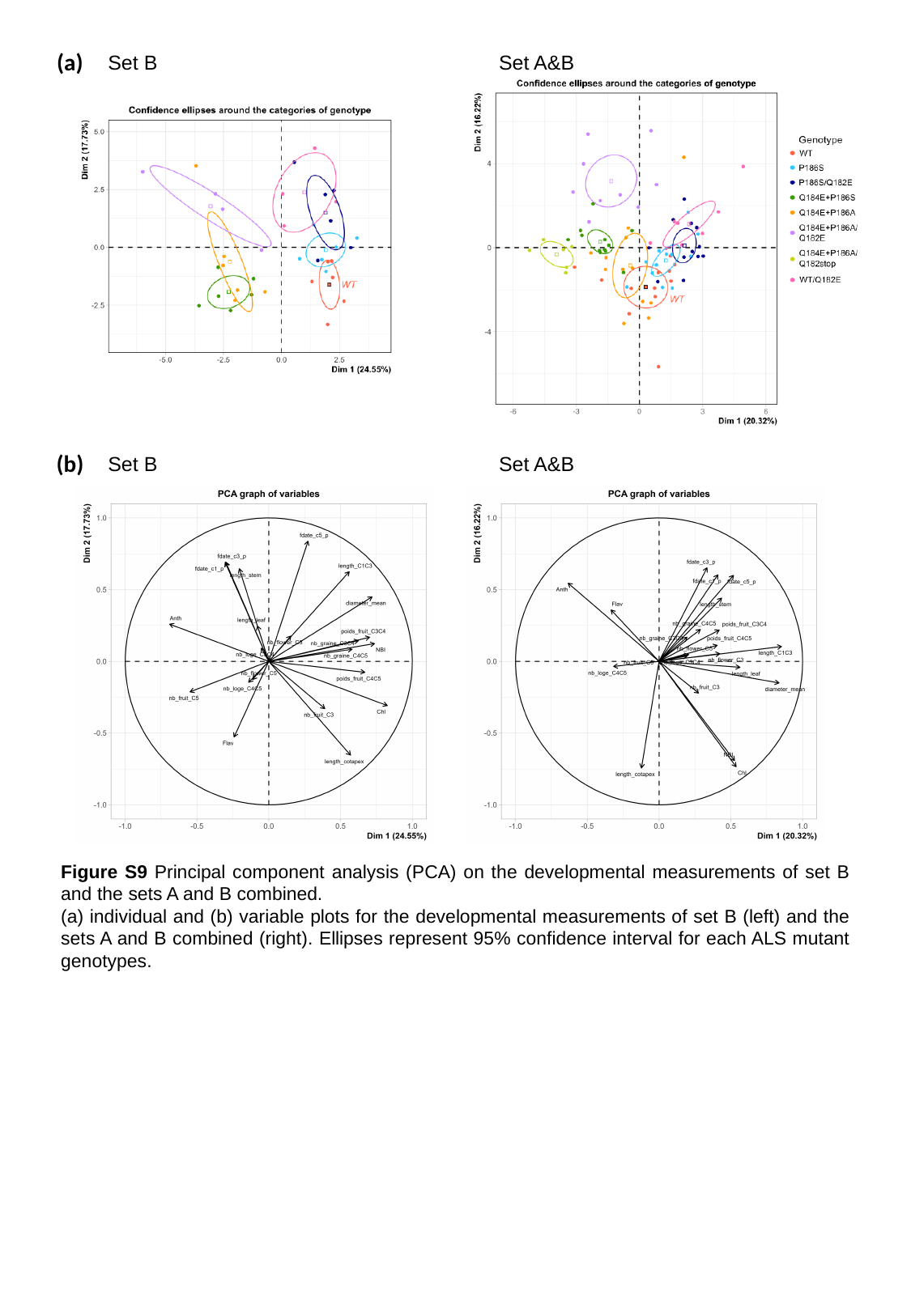

(a)
Set B
Set A&B
(b)
Set B
Set A&B
Figure S9 Principal component analysis (PCA) on the developmental measurements of set B and the sets A and B combined.
(a) individual and (b) variable plots for the developmental measurements of set B (left) and the sets A and B combined (right). Ellipses represent 95% confidence interval for each ALS mutant genotypes.
